# Express barcoding with NextGenPCR and MinION for species-level sorting of ecological samples

**DOI:** 10.1101/2023.04.27.538648

**Authors:** Cristina Vasilita, Vivian Feng, Aslak Kappel Hansen, Emily Hartop, Amrita Srivathsan, Robin Struijk, Rudolf Meier

## Abstract

1. DNA barcodes are useful for species-level sorting of specimen samples, but rarely used in time-sensitive projects that require species richness estimates or identification of pest or invasive species within hours. The main reason is that existing express barcoding workflows are either too expensive or can only be carried out in very well equipped laboratories by highly trained staff.
2. We here introduce a simple workflow that combines rapid DNA extraction with HotSHOT, amplicon production with the aid of NextGenPCR® thermocyclers, and sequencing with low-cost MinION sequencers.
3. We demonstrate the power of the approach by generating and identifying 250 barcodes for 285 specimens within 6 hours. The workflow only required the following major equipment that easily fits onto a lab bench: Conventional thermocycler, NextGenPCR® thermocycler, microplate sealer, Qubit, and MinION.
4. We argue that species-level sorting with simplified barcoding workflows is now faster, more accurate, and sufficiently cost-effective that it can and should replace morphospecies sorting in many projects.

## Introduction

In a perfect world, all biologists should be able to go from a specimen sample to an estimate of species richness and identifications within hours without having to deal with the accuracy problems of morpho-species sorting by parataxonomists or complicated DNA extraction protocols, time-consuming PCR, and access to a capital-extensive sequencing facility. One method that could deliver such fast species-level sorting is a simple express barcoding protocol suitable for time-sensitive ecological projects, pest identification, food authentication, Bioblitzes, and university courses. Express barcoding would also be useful for field stations because researchers could optimize sampling based on samples obtained within hours by newly placed traps and/or initiate the search for immature stages of important species initially only collected as adults.

Developing express barcoding techniques has thus been pursued for almost 15 years. Ivanova et al. (2009) first achieved it for minibarcodes using an optimized workflow that required approximately 2 hours and consisted of the following steps: 5 min DNA extraction of 55 samples (suitable samples: insect fragments, FTA cards and blood), 25 min PCR, 25 min cycle-sequencing, 10 min cleanup, 45 min capillary sequencing, and 5 min trace file analysis. However, the workflow failed to become popular which we surmise was due to the high capital and manpower cost of a laboratory capable of *in situ* high-throughput Sanger sequencing. Such a lab would have to own its own AIT capillary sequencer and pipetting robots run by well-trained lab technicians, because Sanger sequencing requires many repetitive tasks that have to be executed for each specimen. They are DNA extraction, PCR, amplicon clean-up, cycle-sequencing, cycle-sequencing reaction clean-up, capillary sequencing, and trace file bioinformatics.

More recently, alternatives to express barcoding *sensu* Ivanova et al. (2009) have become feasible with the advent of nanopore sequencing developed by Oxford Nanopore Technologies (ONT). ONT sequencers not only have low capital costs, but also require the execution of fewer repetitive tasks, because amplicons can be pooled directly after PCR; i.e., no pipetting robot is needed for tasks such as cycle-sequencing, cleanup, and capillary sequencing. MinION devices are suitable for barcoding specimens under field conditions (Johnson et al., 2017; Parker et al., 2017; Pomerantz et al., 2018; 2022). However, the published methods are not particularly cost-effective because they often require whole ONT flowcells for barcoding few specimens (Maestri et al., 2019; Menegon et al., 2017; Pomerantz et al., 2018), involve expensive and time-consuming molecular procedures, and/or require a complicated bioinformatics procedures. For example, the DNA is extracted using kits requiring tissue digestion, the amplicons are tagged using two PCR reactions (Seah et al., 2020), or the DNA for each sample is quantified before pooling (Maestri et al., 2019). These workflows are particularly impractical when hundreds of specimens need to be processed quickly, which is often the case in ecological studies.

Here, we show that these issues can be overcome by combining DNA “leaching” with HotSHOT, amplicon pooling, MinION sequencing, and the use of fast NextGenPCR® thermocyclers. The latter are still underutilized in molecular ecology although they have been widely used in COVID-19 detection because 30 PCR cycles can be completed within 2 minutes for short DNA fragments. Such fast cycling is achieved through special polymerases and the compression of microplate wells that allow for fast heating/cooling of reagents in three heating zones preset to optimal denaturing, annealing, and extension temperatures. We here show that by combining the fast cycling of NextGenPCR® with MinION sequencing, one can generate and analyze 250 barcodes in 6 hours. All the processes are so robust that they can be learned within days and are suitable for use in university classes. We then suggest that it is time that many projects replace species-level sorting based on morphology with species-level sorting with DNA barcodes (Wang et al., 2018).

## Materials and Methods

Two lab members carried out the experiments that followed Srivathsan et al. (2021) unless mentioned otherwise (see here for videos describing the lab procedures: https://www.youtube.com/@integrativebiodiversitydis5672). We used 10 μl of alkaline lysis buffer for three 96-well plates (285 specimens and 3 control wells) of specimens belonging to Phoridae (Diptera), but the same techniques work for many taxa (e.g., see Yeo et al., 2021). DNA was obtained within 20 min (18 min: 65°C; 2 min: 98°C) and the specimens were recovered intact. Afterwards, 10 μl of neutralising buffer was added to achieve suitable pH (pH 8.1) for DNA storage. For the pipetting of lysis and neutralising buffers manual multichannel pipettes were used. The PCR recipe was as follows: 3 μl of template, 1 μl each of primers (10 μM), 5 μl of the Arctic Fox HF Chemistry-2x (#50050, MBS) polymerase mix. We amplified a 313 bp cytochrome c oxidase subunit I (COI) minibarcode using tagged primers: m1COlintF (Leray et al., 2013) and modified jgHCO2198 (Geller et al., 2013). The PCR plates were sealed with a foil-backed seal using a NextGenPCR® Semi-Automatic Plate Sealer that operates on high temperatures (170°C for two seconds). A resealing step was performed to ensure proper sealing and avoid pre-PCR cross contamination. The PCR reaction was performed on a NextGenPCR® Thermal Cycler (#10001, MBS) and took 22 min with the following conditions: Initial denaturation for 30s at 98°C followed by 5 cycles at 98°C for 10s, 45°C for 30s, 68°C for 10s, followed by 30 cycles at 98°C for 5s, 45°C for 15s and 68°C for 10s.

For the assessment of PCR success rates, agarose gel electrophoresis was performed for 8 samples of Plate 1 and Plate 2 (including the negative controls). Afterwards, the PCR products were pooled (4 μl each) and 100 μl of mixed amplicons was cleaned using 0.8X AMPure XP magnetic beads in a 30 minute wash protocol. All access to the PCR products involved piercing the sealing film with pipette tips mounted on a multichannel pipette. The DNA library was prepared with the Nanopore ligation kit (SQK-LSK114) and NEBNext Companion Module kit (E7180S). Nanopore sequencing (Run 1) of the tagged amplicons employed a flowcell FLO-MIN114 with Oxford Nanopore MinION in a Mk1B and base-calling was conducted using a local computer using Guppy v. 6.3.7+532d62603 and super-accuracy model. Sequencing was stopped after 30 minutes. We resequenced (Run 2) the same pool again using a flowcell dongle “Flongle” FLO-FLG114 because this low-cost flowcell is particularly interesting to ecological labs that only occasionally need a few hundred barcodes. The library preparation used the same procedures, but with the use of less consumables as recommended by the official Flongle library protocol of ONT. Reads were base-called using the super-accuracy model (Guppy v. 6.4.2+97a7f0659). Demultiplexing and barcode calling used ONTbarcoder (2021) under default settings. The filtered barcodes were blasted using SequenceID in GBIF; 95% Query coverage, >89% Identity. Sequences were clustered with Objective Clustering (Meier et al., 2006) at two thresholds; 2% and 4% (Srivathsan et al., 2019).

## Results

We went from 285 specimens to 250 identified barcodes within 6 hours (Figure 1: start: 9:00 am, completion: 3:00 pm) although only one regular thermocycler was used for the HotSHOT DNA extractions and one NextGenPCR® machine for generating amplicons. Agarose gel electrophoresis of the subsamples (8 per plate) from Plates 1 & 2 showed 100% PCR success rates but we only obtained 250 barcodes because of technical handling issues with some PCR reactions in Plate 3 that were traced back to the plate preparation step. All amplicons were sequenced twice. The first time using a fast method by obtaining 507,230 raw reads from a MinION flow cell that only sequenced for 30 minutes (Table 1). The second time, we used a more cost-effective method by employing a Flongle flow cell that produced 321,462 reads in 24 h, generating a total of 241 barcodes (fast method: 250 barcodes). Even with the slow sequencing method, >75% of the barcodes only required the reads obtained within an hour of Flongle sequencing (see Table 1 for results after 1–24 hours).

**Table 1:** Barcoding success rates using MinION and Flongle flow cells.

|  | Number of reads | Plate 1 | Plate 2 | Plate 3 | Overall |
| --- | --- | --- | --- | --- | --- |
| **MinION flow cell** |  |  |  |  |  |
| **0.5h of sequencing** | 507,230 | 97.89% | 91.58% | 73.68% | 87.72% |
| **Flongle flow cell** |  |  |  |  |  |
| **24h of sequencing** | 321,462 | 95.79% | 89.47% | 68.42% | 84.56% |
| **3h of sequencing** | 139,963 | 94.74% | 85.26% | 67.37% | 82.46% |
| **2h of sequencing** | 98,885 | 93.68% | 82.11% | 63.16% | 79.65% |
| **1h of sequencing** | 53,440 | 90.53% | 78.95% | 60.00% | 76.49% |

**Figure 1:**
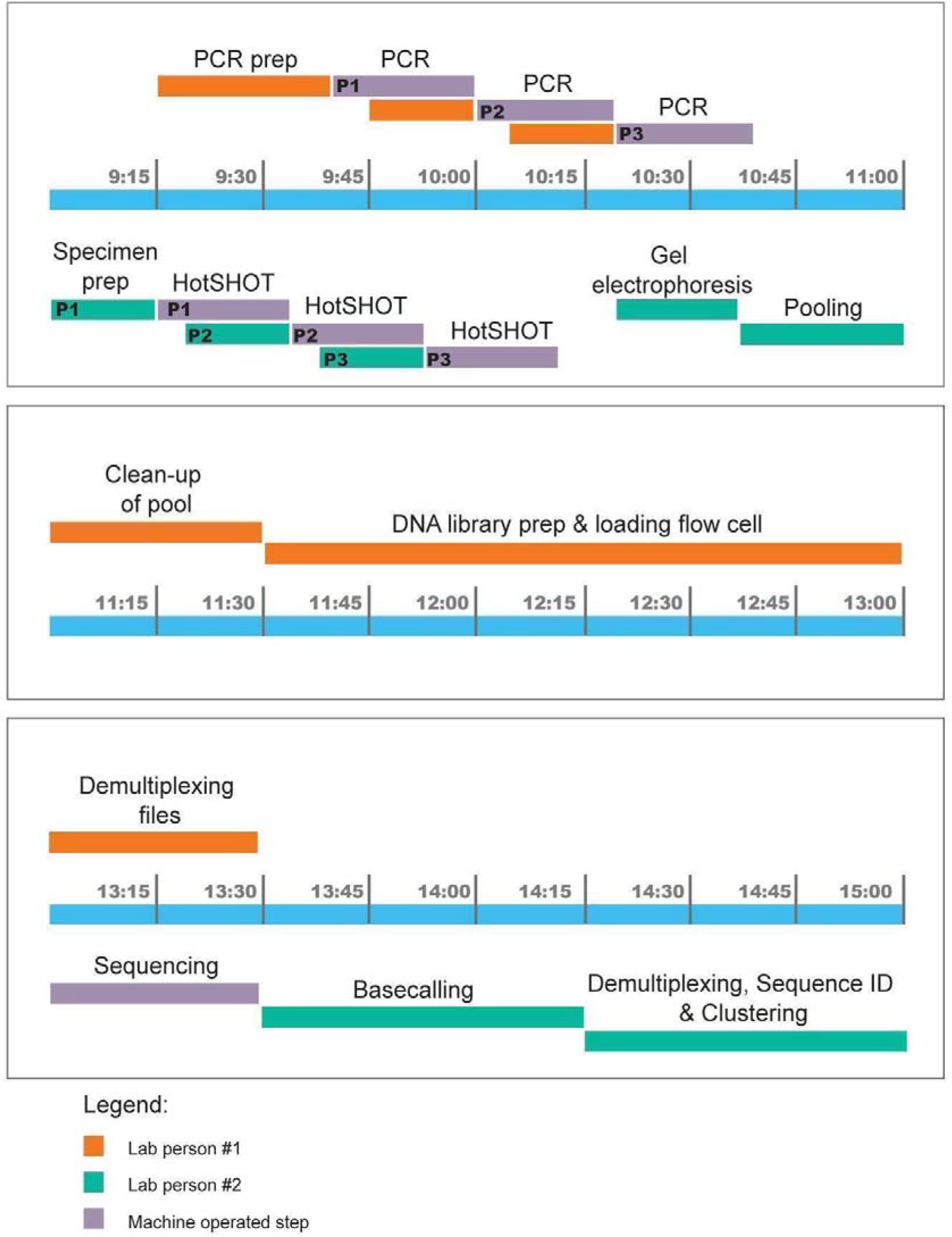
Experiment timeline in blocks of 2 hours (9.00 am – 15.00 pm).

All barcodes were identified by blasting them via the user-friendly “SequenceID” that is implemented on GBIF (https://www.gbif.org/tools/sequence-id: accessed 04 Nov 2022), 249 of the 250 sequences were Phoridae (species belonging to *Anevrina, Megaselia, Phora*, and *Triphleba*). The only non-phorid barcode belonged to *Apis mellifera*, which is known to be parasitised by *Megaselia rufipes* (Dutto and Ferazzi, 2014). Query coverage was over 95% in all sequences; 19 sequences retrieved an exact match, 207 sequences had a close match (90–98.7%) and 24 sequences had a weak match (89.4–89.7%). Objective clustering at multiple thresholds (2–4%) yielded a species richness estimate of 27–30 mOTUs.

## Discussion

Fast species-level sorting of samples consisting of hundreds of specimens is a bread-and-butter task in ecology, university classes, Bioblitzes, etc.. Currently, it involves unpleasant choices because none of the available techniques for species-level sorting are fast, cheap, and accurate. In many projects, species-level sorting is still delegated to parataxonomists who can be quick and cost-effective, but also produce results of unpredictable accuracy because the taxonomic training varies between parataxonomists and the complexity of samples varies between projects and sites (Oliver and Beattie, 1993; Krell, 2004; Baraloto et al., 2007; Derraik et al, 2010; Egli et al., 2020). At the other end of the scale in terms of speed, cost, and accuracy is the identification of specimens by taxonomic experts. In 2006, Marshall et al. provided a cost and time estimate for such expert identification, which was inflation-adjusted USD 3.30 per specimen. On average, the identification took 10.65 minutes, but the 10 taxonomic experts employed by the project could identify only approximately half of the specimens. Presumably, this was due to the lack of taxonomic expertise or identification tools for the remaining specimens.

We here show that species-level sorting with express barcodes can be achieved within hours with a modest amount of equipment, manpower and training. This is achieved by fast cycling using a NextGenPCR® thermal cycler with cost-effective third-generation sequencing with MinION. Barcoding required only six hours using techniques that can be learned within days. This distinguishes the new workflow from other express barcoding methods that have been available for many years but are rarely used due to high complexity, capital, manpower, and/or consumable cost. We here show that express barcoding only requires one NextGenPCR® thermal cycler (including plate sealer) in addition to standard equipment found in most molecular laboratories (pipettes, Qubit, regular thermocycler, MinION). Like all other express barcoding protocols, we achieved short PCR cycling times in part by using special enzymes that, however, are responsible for much of the cost. Ivanova et al. (2009) recommended TaKaRa Z-Taq, or KAPA 2G Fast, while we here used NextGenPCR® Arctic Fox HF which costs ca. €1.70 c.q. $1.85 USD per specimen. For those ecologists who do not need barcodes within 24 hours, costs can drop to below USD 1 using modifications discussed below; i.e., barcoding can be already be cheaper than expert identification (Marshall et al., 2006).

We here generated 250 barcodes for 285 specimens in 6 hours using a partial MinION flow cell and two staff members (ca. 2.5 minutes/specimen). The success rate (87.72%) would have been even higher if we had not observed unusual evaporation in the wells of one row of the third plate, which was probably not properly sealed. Overall, this means that express barcoding does not have to mean lower barcoding success rates than what is obtained with regular protocols. In addition, the experiment could have been completed even faster if we had used three parallel thermal cycling setups instead of only one regular thermocycler for DNA extraction and one NextGenPCR® thermocycler for PCR. We estimate that the use of three cyclers would have reduced the processing time by another hour. However, we here wanted to report realistic times for those users (e.g., field station, citizen science project) that would like to implement express barcoding with a minimal amount of equipment. We find that such realistic estimates were hard to obtain for other express barcoding protocols because they were, for example, vague about the time needed for DNA extraction, tagging amplicons (often done using two PCR reactions), and specimen handling. Note that the times are for 313 bp minibarcodes obtained for phorid flies, but we have used similar protocols for many other insect taxa (Lee et al., 2022; Yeo et al., 2018; Yeo et al., 2021; Ho et al., 2020; Srivathsan et al., 2022). Furthermore, we have shown that the use of minibarcodes yields similar identification success rates as the use of full-length barcodes (Yeo et al., 2020).

Two protocol variations are attractive to users who prioritize low cost over speed. The first is the use of a Flongle instead of a MinION flow cell for sequencing. This adds several hours to protocol because Flongle flow cells have fewer pores and thus sequence more slowly. The delay depends on the targeted success rate and the number of amplicons in the pool (Table 1). If only 200 specimens have to be barcoded, we show that 3 hours of Flongle sequencing will yield similar success rates as 15–20 minutes of MinION flow cell sequencing. The attractive aspect of using a Flongle is the low consumable cost of only ca. USD 120 (including library prep). Obtaining the same data with a standard MinION flow cell would be more expensive, but the precise cost is hard to estimate because regular MinION flow cells can be washed and reused. A second cost-saving modification of the express barcoding workflow presented here involves the use of cheaper polymerases and several traditional thermocyclers. For example, when using three thermocyclers simultaneously, the DNA in the three microplates could be amplified using conventional enzymes (a regular high performance enzyme USD 0.60 per specimen or the extremely cheap options for USD 0.10 per specimen) within two hours instead of the total one hour needed with a single NextGenPCR® cycler. However, the use of more equipment can be unattractive for field stations, as space and/or electrical outlets are at a premium. Longer waiting times are also unattractive when members of the public in citizen science projects or students in university classes are waiting to see the results. However, using a Flongle and conventional thermocyclers means that the cost per barcode drops well below USD 1 per specimen; i.e. barcoding will be significantly cheaper than expert sorting/identification of specimens (Marshall et al., 2006). Note, however, that the latter can yield more species names, because barcode databases still lack identified sequences for many species.

## Acknowledgements

AKH acknowledges the Carlsberg Foundation for their continuous support of his postdoc activities through the project “Next Generation Taxonomy”. CV was supported by the Bundesministerium für Bildung und Forschung, Berlin, Germany, project “German Barcode of Life III: Dark Taxa” (FKZ 16LI1901C). The producer of NextGenPCR instruments and reagents, Molecular Biology Systems, provided the reagents and 7Bioscience GmbH access to a NextGenPCR® thermocycler.

## Author contributions

Cristina Vasilita and Rudolf Meier conceived and designed the project; Cristina Vasilita, Vivian Feng and Amrita Srivathsan collected the data; Cristina Vasilita, Vivian Feng and Amrita Srivathsan analysed the data; Emily Hartop and Aslak Kappel Hansen contributed valuable discussion of the obtained results; Robin Struijk optimized barcoding protocols for NextGenPCR® thermocyclers and coordinated access. All authors contributed to the manuscript and gave final approval for submission.

